# Prediction of potential disease-associated microRNAs using structural perturbation method

**DOI:** 10.1101/223693

**Authors:** Xiangxiang Zeng, Li Liu, Linyuan Lü, Quan Zou

## Abstract

**Motivation:** The identification of disease-related microRNAs(miRNAs) is an essential but challenging task in bioinformatics research. Similarity-based link prediction methods are often used to predict potential associations between miRNAs and diseases. In these methods, all unobserved associations are ranked by their similarity scores. Higher score indicates higher probability of existence. However, most previous studies mainly focus on designing advanced methods to improve the prediction accuracy while neglect to investigate the link predictability of the networks that present the miRNAs and diseases associations. In this work, we construct a bilayer network by integrating the miRNA–disease network, the miRNA similarity network and the disease similarity network. We use structural consistency as an indicator to estimate the link predictability of the related networks. On the basis of the indicator, a derivative algorithm, called structural perturbation method (SPM), is applied to predict potential associations between miRNAs and diseases.

**Results:** The link predictability of bilayer network is higher than that of miRNA–disease network, indicating that the prediction of potential miRNAs-diseases associations on bilayer network can achieve higher accuracy than based merely on the miRNA–disease network. A comparison between the SPM and other algorithms reveals the reliable performance of SPM which performed well in a 5-fold cross-validation. We test fifteen networks. The *AUC* values of SPM are higher than some well-known methods, indicating that SPM could serve as a useful computational method for improving the identification accuracy of miRNA-disease associations. Moreover, in a case study on breast neoplasm, 80% of the top-20 predicted miRNAs have been manually confirmed by previous experimental studies.

**Availability and Implementation:** https://github.com/lecea/SPM-code.git

**Supplementary information:** Supplementary data are available at *Bioinformatics* online.

## 1 Introduction

MicroRNAs (miRNAs) are a class of small, endogenous, non-coding RNAs that function in post-transcriptional regulation of gene expression and RNA silencing (Ambros, 2004; Bartel, 2004). Accumulated experimental evidence suggests that miRNAs are involved in close relationships with the emergence and development of human diseases (Alvarez-Garcia and Miska, 2005; Jopling, *et al.*, 2005). Therefore, identifying the associations between miRNAs and diseases may largely help to understand the diseases’ pathogeneses. Multiple databases, such as HMDD v2.0 (Li, *et al*, 2014), miR2Disease (Jiang, *et al.*, 2009), miRCancer (Xie, *et al.*, 2013), and dbDEMC (Yang, 2010) have been constructed to store useful data regarding RNAs.

Given that biological experimental methods involve many restrictions, such as high costs and long execution times, computational prediction methods are useful by prioritizing candidate miRNAs for specific diseases (Zeng, *et al.*, 2015). How to well utilize the known information to predict the potential disease-associated miRNAs? The question is actually the link prediction problem on miRNA–disease networks (Lü, *et al.*, 2011). Jiang, *et al.* (2010a) proposed the first computational method, which constructed a human phenome-miRNAome network. The similarity scores were calculated by the cumulative hypergeometric distribution. To further ameliorate their method, Jiang, *et al.* (2010b) integrated multiple genomic data using the naïve Bayes model. The scholars then calculated the functional similarity between genes under the biological knowledge that miRNAs regulate diseases through their target genes. Xu, *et al.* (2011) focused on extracting features from the miRNA–disease network data. These network data were constructed under two considerations, namely, a feature set primarily related to miRNA information and disease phenotype information. Chen, *et al.* (2012) presented the RWRMDA model to identify potential miRNA–disease pairs by adopting random walks on the miRNA functional similarity network. Moreover, Shi, *et al.* (2013) improved the RWRMDA by considering miRNA–target associations, disease–gene associations, and protein–protein interaction networks. HDMP (Xuan, *et al.*, 2013) evaluated the *k* most functionally similar neighbors by considering the disease terms and phenotype similarity. Chen and Yan (2014) presented a semi-supervised method based on regularized least squares (RLSMDA). In the study, the authors used their proposed method to integrate known miRNA–disease associations, disease–disease similarity datasets, and miRNA–miRNA functional similarity networks. Xuan, *et al.* (2015) developed the MIDP method, which uses the features of different nodes on the basis of random walks with a restart. An extension method, named MIDPE, was proposed by constructing a miRNA–disease bilayer network. This approach was formulated because nearly all the previous methods based on random walks could not be applied without any known related miRNA. The KATZ method (Zou, *et al.* 2015; Lü, *et al.*, 2009) denotes the associations on the basis of paths of different lengths in the miRNA–disease network. (Luo, et al., 2016) used the local neighbors of different node types and developed a collection prediction method on the basis of transduction learning (CPTL) to predict the miRNA–disease interactions. WBSMDA method (Chen, *et al.*, 2016) integrates several heterogeneous biological datasets on the basis of between-scores and within-scores for miRNA–disease associations. Meanwhile, PBMDA (You, *et al.*, 2017) is an effective path-based model for miRNA–disease association prediction. This model adopts the depth-first search algorithm by integrating the disease semantic similarity, miRNA functional similarity, known human miRNA–disease associations, and Gaussian interaction profile kernel similarity for miRNAs and diseases.

The existing methods can be categorized into four aspects: (i) neighborhood-based methods, such as HDMP (Xuan, *et al.*, 2013) and CPTL (Luo, *et al.*, 2016); (ii) random walk-based methods, such as RWRMDA (Chen, *et al.*, 2012), Shi’s method (Shi, *et al.*, 2013), MIDP, and MIDPE (Xuan, *et al.*, 2015); (iii) machine learning-based methods, such as Jiang’s method (Jiang, *et al.*, 2010), Xu’s method (Xu, *et al.*, 2011), and the RLSMDA (Chen and Yan, 2014); (iv) path-based methods, such as KATZ (Zou, *et al.*, 2015) and PBMDA (You, *et al.*, 2017).

Most previous studies mainly focus on designing advanced methods to improve the prediction accuracy while neglect to investigate to what extend the miRNAs–diseases associations can be predicted. In this study, we use an indicator for estimating link predictability, named the structural consistency index (Lü, *et al.*, 2015), to solve the above problems. Given the perturbation of the adjacency matrix, the structural consistency index is free of prior knowledge of network organization. The predictability of a network is reflected by the consistency of the structural features before and after a removal of a part of links randomly. Then, we apply the structural perturbation method (SPM), to predict potential miRNA-disease associations. In the prediction process, we first construct a bilayer network which integrates the verified miRNA–associated diseases with the disease and miRNA similarity networks. Then, we randomly select a part of links from the bilayer network to form a perturbation set. We use the perturbation set to agitate the remaining links by first-order approximation and then compute the perturbed adjacency matrix. The unobserved links are ranked by their scores in the perturbed matrix. The top-ranked miRNAs are selected as the prediction results.

We summarize our major contributions as follows. First, a bilayer network is constructed for miRNA–disease prediction. This network integrates three single networks, namely the miRNA□miRNA similarity network, the disease disease similarity network, and the miRNA disease bipartite network where the edges represent the associations between miRNAs and diseases. Second, we employ the structural consistency indicator to investigate the link predictability of the constructed bilayer network and find that it is predictable with structural consistency equal to 0.581 which is much higher than any of the three single networks. Third, we apply the SPM to predict potential miRNA-disease associations. We select new associations that cannot seriously affect the structural consistency of a network. Finally, we execute an iterative operation for effectively improving the performance, given that the miRNA-disease bilayer network is incomplete. The experimental results show that our method can achieve higher accuracy than many previous methods in a 5-fold cross-validation. Moreover, in a case study on breast neoplasm, 80% of the top-20 predicted miRNAs have been manually confirmed by previous experimental studies.

## 2 Methods

### 2.1 Construct miRNADdisease bilayer network

We first construct a bilayer network which integrates three networks, namely the bipartite network with verified miRNA-disease associations, the diseases similarity network, and the miRNAs similarity network.

To construct the miRNA–disease network, we download the latest version of the human miRNAD disease database (HMDDv2.0) (Li, *et al.*, 2014). If a miRNA is related to a disease, an edge is added to link them. A total of 6,441 associations between 577 miRNAs and 336 diseases are available after removing duplications.

Functionally similar genes demonstrate a greater probability of regulating similar diseases. Therefore, we use gene functional information to construct disease similarity network. We download the data from the HumanNet database (Lee, *et al.*, 2011), which contains an associated log-likelihood score *(LLS)* of each interaction between two genes or gene sets. We calculate the similarity between diseases and based on the gene functional information, as follows:

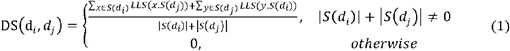

where *S*(*d_i_*) and *S*(*d_j_*) represent the gene sets that related to diseases and *d_j_*, respectively. |*S*(*d_i_*)| and |*S*(*d_j_*)| are the cardinalities of gene sets *S*(*d_i_*) and *S*(*d_j_*), respectively. *LLS*(*x*, *S*(*d_j_*)) is the *LLS* between gene *x* and gene set *S*(*d_j_*) where *x* ≈ *S*(*d_i_*). Similarly, we can define *LLS*(*y*, *S*(*d_i_*)). If *DS*(*d_i_*, *d_j_*), it can be considered as the weight of the link connecting diseases and *d_j_*. Hence, we obtain a weighted disease similarity network containing 112,896 similar associations between 336 diseases.

The miRNA similarity network is constructed by employing four main parameters of two miRNAs, namely the verified miRNA–target associations, family information, cluster information, and verified miRNA disease associations. First, we download the verified miRNA–target associations experimentally supported from the miRTarBase (Hsu, *et al.*, 2014). Two miRNA nodes are connected if they share common targets; the edge weight, *RST* (miRNA Similarity based on Target), represents the number of shared targets between miRNAs. Therefore, we obtain 463,483 pairwise *RST.* We subsequently download the miRNA family data from ftp://mirbase.org/pub/mirbase/CURRENT/mi-Fam.dat.gz (Griffithsjones, *et al.*, 2003). Different miRNAs belong to 299 miRNA families. If two miRNAs belong to the same miRNA family, we set their *RSF* (miRNA Similarity based on Family) value as 1, otherwise 0. Third, the miRNA cluster information is accessible in miRbase (Kozomara and Griffithsjones, 2014). We obtain 153 clusters of miRNAs. If two miRNAs belong to the same cluster, then the *RSC* (miRNA Similarity based on Cluster) value set as 1. Obviously, *RSF* and *RSC* are both Boolean type matrix. Finally, according to pervious literatures (Wang, *et al*., 2010), miRNAs with similar function are more likely to associate with similar diseases. Therefore, we utilize MISIM (downloaded from http://www.cuilab.cn/files/images/cuilab/misim.zip), a miRNA similarity network based on verified miRNA–disease associations, to define *RSD* (miRNA Similarity based on Disease). After data preparation, we combine the *RST, RSF, RSC*, and *RSD* to calculate the similarity *RS*(*r_i_*, *r_j_*) between miRNA *r_i_* and miRNA *r_j_*, as follows:

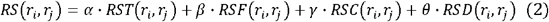

where *α*, *β*, *γ* and *θ* are the corresponding parameters to adjust the weights of the four parts, and *α* + *β* + *γ* + *θ* = 1. *RST, RSF, RSC* and *RSD* values are calculated based on the miRNA–target, miRNA family, miRNA cluster, and miRNA–disease, respectively. Selecting the suitable values for *α*, *β*, *γ* and *θ* is essential. In our experiments, we tune the parameters *α*, *β*, *γ* and *θ* from 0 to 1 with step 0.1, whereas *θ*= 1 − *α* − *β* − *γ*. The prediction achieves the best performance when *α* = 0.2, *β* = 0.1, *γ* = 0.2 and *θ* = 0.5. The *RS* value between two miRNAs can be considered as the weight of the link if it is greater than 0. Hence, we obtain a weighted miRNA similarity network which contains 332,928 similar associations between 577 miRNAs.

By integrating the above three networks, we constructed a bilayer network. We define matrix *RSnet* as the miRNA similarity network, the weights are *RS* values. We define matrix *DSnet* as the disease similarity network, the weights are *DS* values. We define matrix *RDnet* as the known disease-miRNA associations, if miRNA *i* and disease *j* are connected, the element *RDnet_ij_* = 1; otherwise, *RDnet_ij_* = 0. Therefore, miRNADdisease bilayer network can be expressed by a *N* × *N*(*N* =913, the total number of miRNAs and disease in the network) undirected adjacency matrix *A_N×N_*.

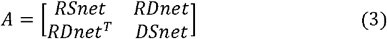

For prioritizing the most possibly potential miRNAD□disease associations, Figure 1 demonstrates a flowchart of how to construct the bilayer network.

**Fig. 1.**
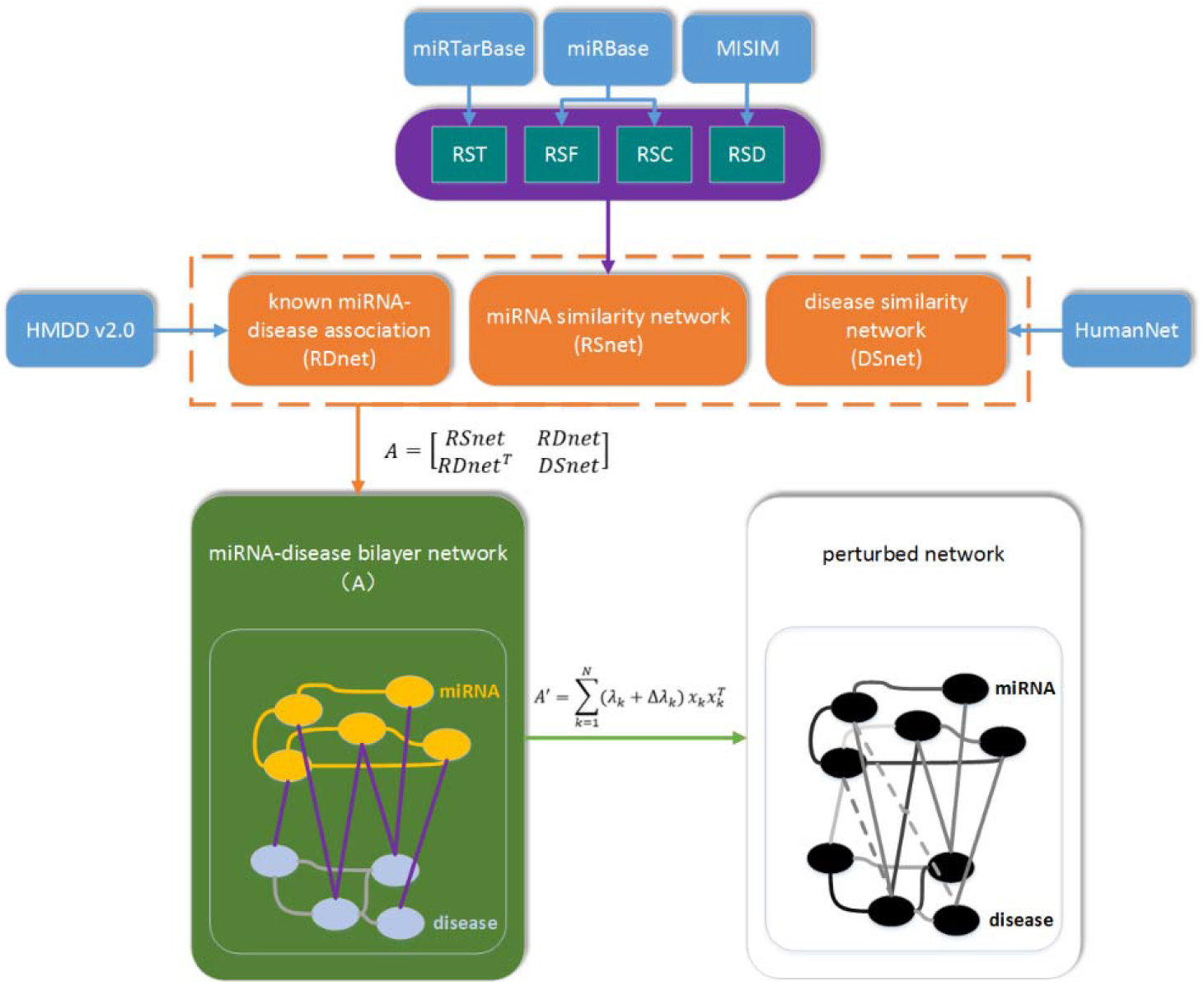
The process of constructing a bilayer network and calculating perturbed network

### 2.2 Structural consistency index

Structural consistency is used to quantify the link predictability of complex network (Lü, *et al.*, 2015). It is defined as the consistency of structural features before and after a removal of a partial of associations randomly. In this paper, we extend this method to weighted bilayer networks.

The weighted miRNAD□disease bilayer network can be presented by a graph *G(V, E, W). V*is the set of nodes, which include both miRNA and disease nodes, *E* is the set of edges, and *W*is the set of weights. We select a fraction of the links to compose a perturbation set Δ*E*, whereas the rest of the links define as *E^R^*. Δ*A* and *A^R^* denote the corresponding weighted adjacency matrices respectively; and *A* = *A^R^* + *Δ^A^*. Obviously, *A^R^* is a real symmetric matrix, therefore *A^R^* can be diagonalized as follows.

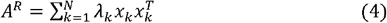

where *λ*_*k*_ are the eigenvalues for, and are the corresponding orthogonal and normalized eigenvectors. Using ΔE as perturbation set, we obtain a perturbed matrix by first-order approximation. First-order approximation allows the eigenvalues to change but keep the eigenvectors constant. Two cases are considered. First, considering the non-degenerated case without any repeated eigenvalues. After perturbation, the eigenvalue *λ_k_* is adjusted to be *λ_k_* + Δ*λ_k_*, and the corresponding eigenvector is adjusted to be *x_k_* + Δ*x*. By multiplying the eigenfunction, we have

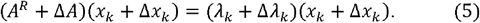

By neglecting the second-order terms 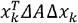 and 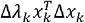, we obtain

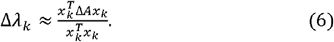

Keeping eigenvectors unchanged, using the perturbed eigenvalues, the following perturbed matrix can be obtained,

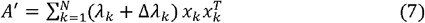

which is considered as linear approximation.

Next, considering the adjacency matrix contains repeated eigenvalues. If *λ_ki_*. as eigenvalues, the index *i* denotes *M* related eigenvectors of the same eigenvalues and the index *k* denotes different eigenvalues. Given that any linear combination of eigenvectors belonging to the same eigenvalue is still an eigenvector. After adding a perturbation into the network, we choose the degenerate eigenvalues, which can be changed successively into the perturbed non-degenerate eigenvalues. If we define the chosen eigenvectors to be 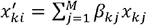 the eigenfunction becomes

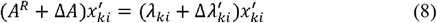

giving us

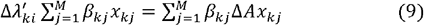

For any *n*= 1 ⋯*M*, left multiplying equation (9) by 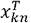, we obtain

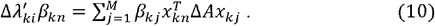

Written in matrix form, equation (10) becomes

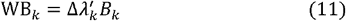

where 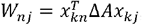 which is a *M×M* matrix, and *B_k_* is the column vector of *β_kj_*. Then, according to eigenfunction (11), we obtain 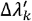 and B_*k*_, the perturbed adjacency matrix *A*′ is obtained by simply replacing *x_k_* and Δ*λ_k_* in equation (7) with 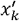 and 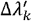.

Matrix eigenvectors can reflect the network structural features. If the eigenvectors of the matrix *A*′ and the matrix *A* are nearly the same, it indicates that the perturbation set not significantly changes the structural features. In other words, the network is of high structural consistency. All unobserved links and perturbed links are ranked in descending order according to their corresponding scores in perturbed matrix *A*′. Denote the set of top-*L* links as *E^L^*, where *L* = |Δ*E*|. Structural consistency σ_c_ is defined as the ratio of shared links between Δ*E* and *E^L^* to *L*, follow as

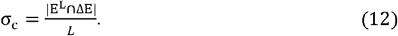

Figure 2 shows how to calculate the structural consistency σ_c_ of an exampled network. The left figure shows the adjacency matrix *A*, where the number in each square is the corresponding value of the matrix element. The second figure represents the matrix *A^R^* which is obtained by randomly removing a fraction of the observed links. The removed links, namely (2,10), (2,14), (3,4), (3,7) and (8,12), constitute the perturbation set Δ*E*. Obviously, *L* = |Δ*E*| = 5, The right figure is the perturbed matrix *A*′. By ranking the unobserved links in according to their corresponding values in *A^R^*, we obtain that the top-*L* links in *E^L^* are (2,10), (3,4), (6,15), (8,12), and (12,15). Therefore, there are three shared links between Δ*E* and *E^L^*, σ_c_ = 0.6.

**Fig. 2.**
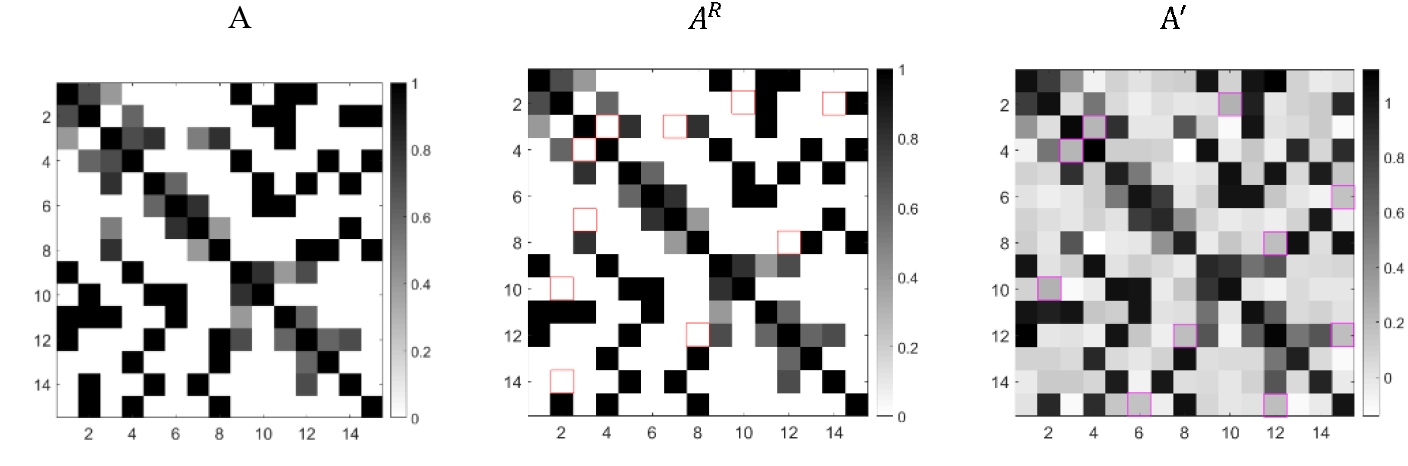
An example of calculating σ_c_

We calculated the structural consistency of the seven related networks mentioned in section 2.1, which is shown in Table 1. For every network, 10% of total links are selected at random to compose the perturbation set. Note that in this experiment, we randomly select links in the bilayer network, which means the perturbation set may contain three kinds of links. The network *RSnet* that contains more information has higher structural consistency than MISIM. The constructed miRNA–disease bilayer network has the highest structural consistency, indicating that higher structural consistency can be obtained by considering more information. Moreover, *RDnet+RSnet* has higher *σ_c_* than *RDnet+DSnet*, indicating that *RSnet* is more helpful than *DSnet* on improving the link predictability.

**Table 1.** The structural consistency of the seven related networks

| Network | MISIM* | RSnet | DSnet | RDnet | RDnet+DSnet | RDnet+RSnet | Bilayer network (A) |
| --- | --- | --- | --- | --- | --- | --- | --- |
| structural consistency | 0.317±0.03 | 0.340±0.06 | 0.316±0.05 | 0.223±0.04 | 0.232±0.01 | 0.275±0.03 | **0.581±0.02** |

\* MISIM is a miRNA similarity network based on verified miRNA-disease associations, and we call it as RSD (miRNA similarity based on disease);

### 2.3 Structural perturbation link prediction method

Different from the experiments in (Lü, *et al.*, 2015), in our paper the perturbation is carried out on a weighted bilayer network. To evaluate the performance of SPM, we adopted 5-fold cross-validation based on verified miRNA–disease associations downloaded from HMDD database (Li, *et al*, 2014). In 5-fold cross-validation, the original set is random divided into five equal sized subsets. Of the five subsets, a single subset is retained as the validation data for testing the model, and the remaining four subsets are used as training data. The cross-validation process is then repeated five times, with each of the five subsets used exactly once as the validation data. The five results from the folds can then be averaged to produce a single estimation. For a bilayer network *A*, we predict miRNA disease potential associations by using SPM. The observed miRNA disease associations are partitioned into five equal subsets randomly. One of the five subsets is selected as probe set, and the rest four subsets together with the *RSnet* and *DSnet* constitute the training set. Then, we randomly remove a fraction of links from training sets to constitute the perturbation set. The perturbed matrix can be obtained as through equation (7) (see section 2.2).

The final prediction matrix 〈*A*′〉 is obtained by averaging over ***t*** independent selections of perturbation set. Under this framework, the entries for 〈*A*′〉 can be considered as score between a pair of nodes of the given bilayer network *A*. Their scores in 〈*A*′〉 determine the rank of all unob-served miRNA–disease associations with the assumption that links with higher scores have greater existence likelihood.

## 3 Results

### 3.1 Evaluation metrics

We evaluate the ability of SPM to predict potential disease-related miRNAs by performing a 5-fold cross-validation. For a specific disease *d*, *d*-related relationships are randomly divided into five subsets, four of which are used as known information; the last subset is used for testing. We introduce three evaluation metrics that are proposed for the evaluation of the performance of miRNAD disease association prediction results.

*PRE* is the *precision* of specific diseases. *PRE* is the ratio of related samples selected to the number of samples selected.

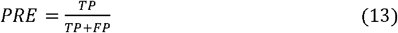

where *TP* and *FP* are the number of true positive and false positive samples with respect to a specific disease, respectively. Based on the definition, the larger *PRE* value, the better prediction accuracy.

*REC* is the *recall* of the specific diseases.

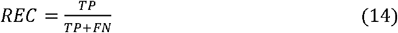

where *FN* is the number of false negative samples with respect to a specific disease.

*AUC* means the area under the receiver operating characteristic curve, which is a global prediction performance indicator. We calculate the true positive rate (*TPR*) and the false positive rate (*FPR*) by varying the threshold, and obtain the receiver operating characteristic (ROC) curves.

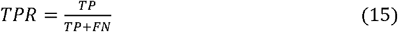

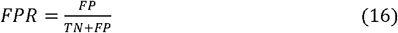

where *FN* and *FP* are the number of negative and positive samples erroneously identified. *TN* and *TP* are the number of negative and positive samples correctly identified.

### 3.2 Performance on the bilayer network

We firstly investigate the effect of parameter *t* on the prediction performance of SPM on bilayer network constructed in Section 2.1. Figure 3 shows the change in *AUC* values as *t* increases. Each point is the average over five *AUC* values. As can been seen that the *AUC* values increase with *t*, but when *t* over ten, the *AUC* values nearly remained unchanged. Therefore, for simplicity we set *t* = 10 for all experiments.

**Fig. 3.**
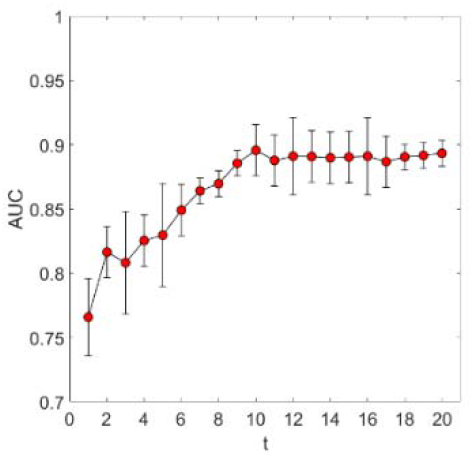
*AUC*values *vs.* parameter *t*

Table 2 shows the *AUC* across all tested diseases of the four related networks. The prediction accuracy of miRNA disease relations on bilayer network is the highest among the four cases, indicating that the addition of *RSnet* and *DSnet* can improve the prediction accuracy.

**Table 2.**
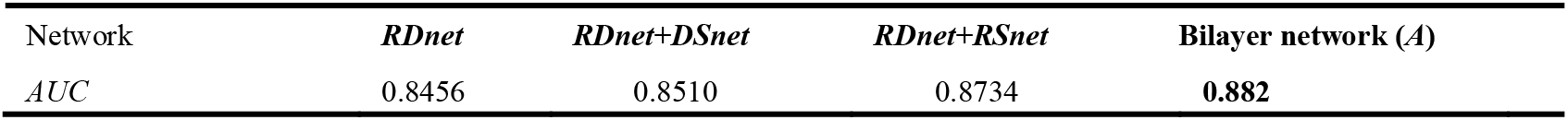
The *AUC* across all tested diseases of the four related networks

The existing literatures indicate that many of diseases are related with only few miRNAs, the number of miRNAs was insufficient to evaluate the prediction performance. Therefore, we tested the performance of SPM in three diseases, namely breast neoplasm, heart failure and lung neoplasm. For a specific disease, we rank the related candidates according to their scores in *A*′. We measured the *PRE* and *REC* within the top 10, 20,…, 90, and 100 candidates in rank list, because the top portion of the prediction links are more important. The *PRE* indicates the ratio of positive samples in the top-k samples. The *REC* measures how many positive samples are correctly identified within the top-*k*. ROC curves were used as the global prediction performance indicator.

Table 3 shows the ROC curves, *PRE*, and *REC* for different diseases. The *AUC* values of the three diseases achieve relative good performance. As *k* increasing, *REC* is on the rise, but *PRE* decreases. This demonstrates that the links ranked the top places have a greater probability of being potential associations.

**Table 3.**
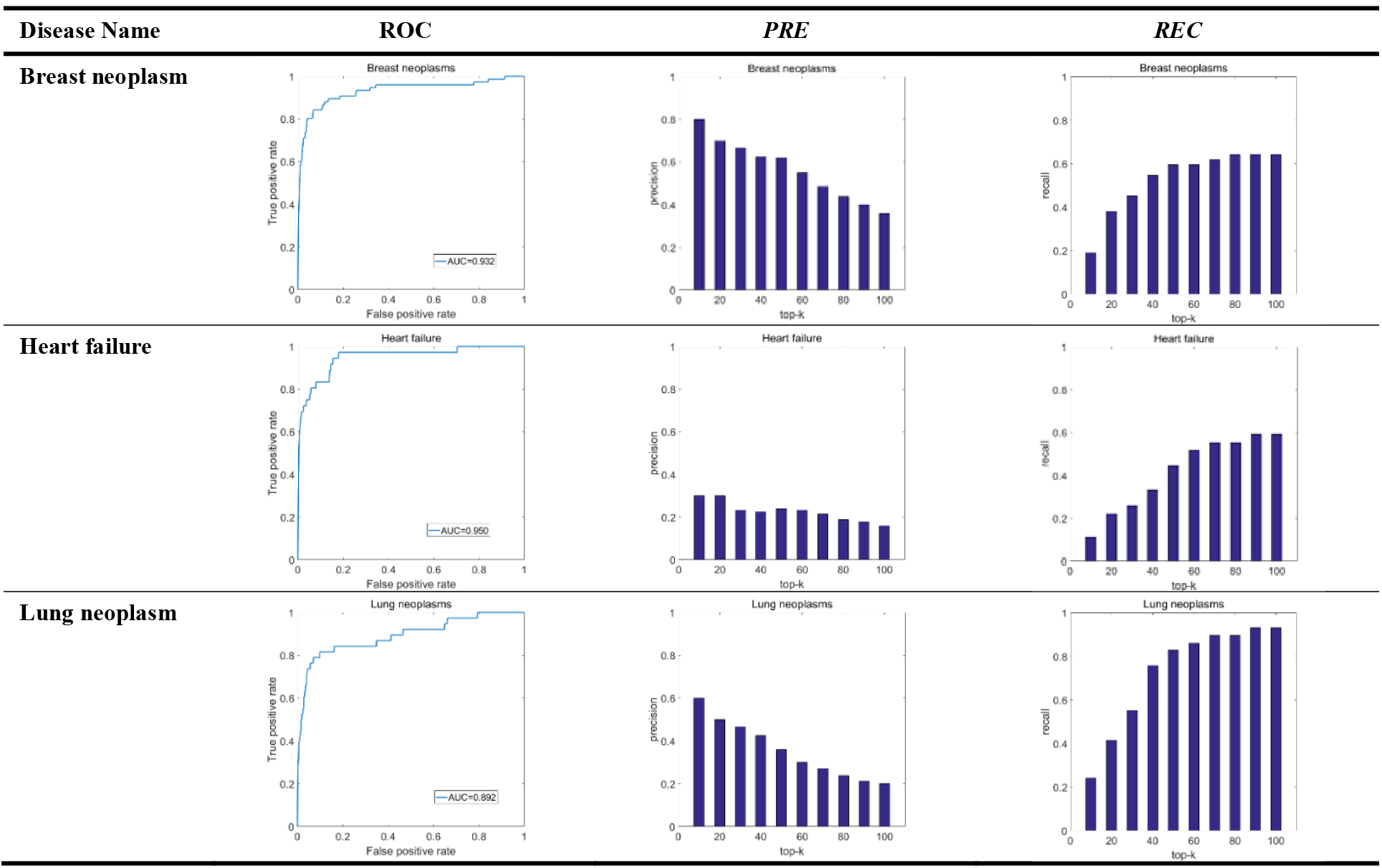
ROC curves, *precision*, and *recall* for SPM by using a 5-fold cross-validation

### 3.3 Comparison with other existing methods

We compare the SPM with four widely applied miRNA-disease prediction algorithms: (i) RWRMDA (Chen, *et al.*, 2012); (ii) HDMP (Xuan, *et al.*, 2013); (iii) RLSMDA (Chen and Yan, 2014); (iv) MIDP (Xuan, *et al.*, 2015). We use the same bilayer network (constructed in section 2.1) for different methods. Table 4 shows the prediction performance measured by *AUC* for different diseases. The highest value for each row is in boldface. For all fifteen tested diseases, SPM is superior to all other algorithms, including MIDP. The average *AUC* values of RWRMDA, HDMP, RLSMDA, MIDP, and SPM are 79.9%, 81.6%, 82.6%, 86.2%, and 91.4%, respectively. The average *AUC* of SPM is higher than that of the other four methods by 11.5%, 9.8%, 8.8%, and 5.2%, respectively. The highest *AUC* value of acute myeloid leukemia is 0.957.

**Table 4.**
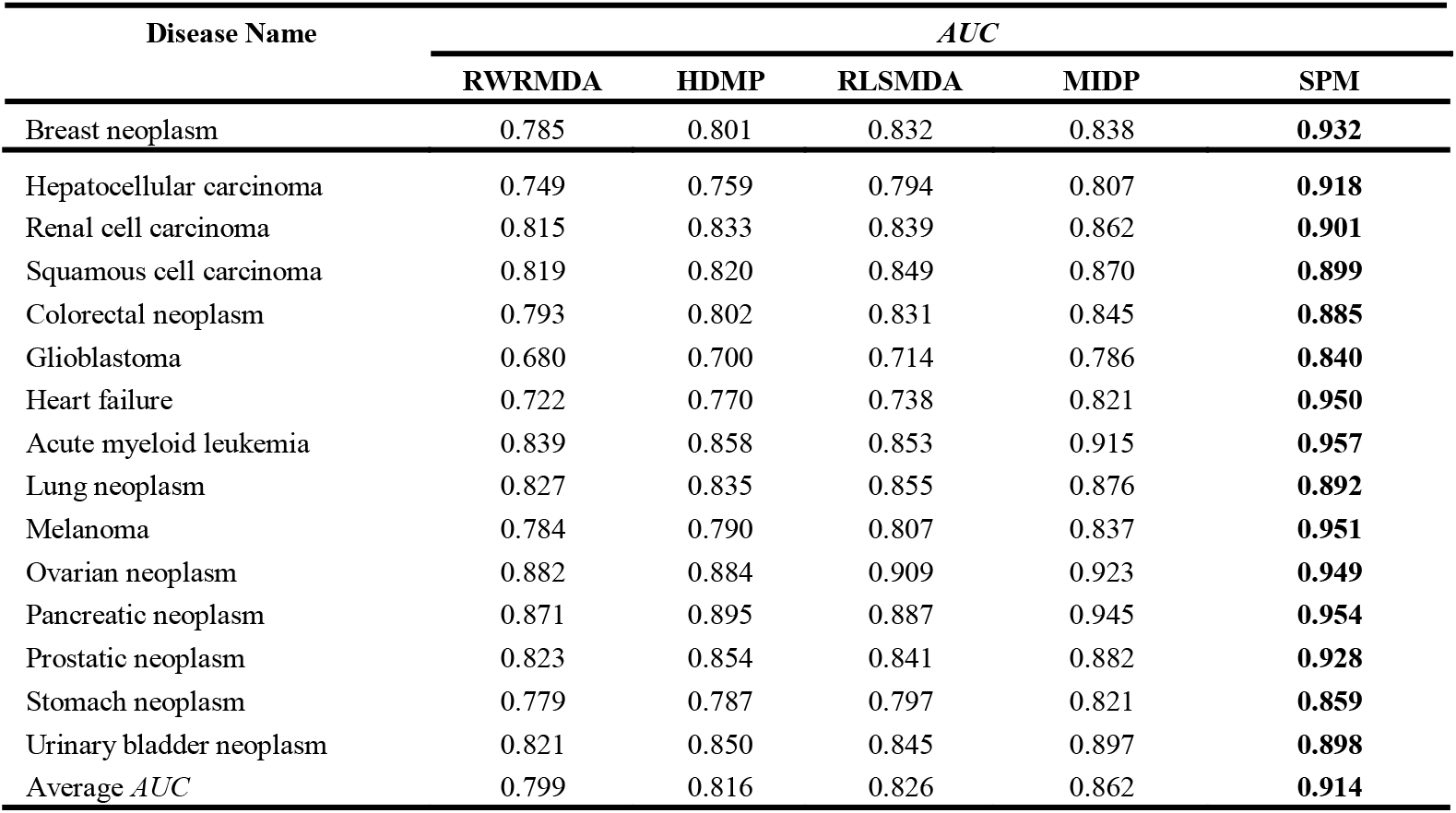
The *AUC* values for five methods by using 5-fold cross-validation

| Disease Name | *AUC* |  |  |  |  |
| --- | --- | --- | --- | --- | --- |
|  | RWRMDA | HDMP | RLSMDA | MIDP | SPM |
| Breast neoplasm | 0.785 | 0.801 | 0.832 | 0.838 | **0.932** |
| Hepatocellular carcinoma | 0.749 | 0.759 | 0.794 | 0.807 | **0.918** |
| Renal cell carcinoma | 0.815 | 0.833 | 0.839 | 0.862 | **0.901** |
| Squamous cell carcinoma | 0.819 | 0.820 | 0.849 | 0.870 | **0.899** |
| Colorectal neoplasm | 0.793 | 0.802 | 0.831 | 0.845 | **0.885** |
| Glioblastoma | 0.680 | 0.700 | 0.714 | 0.786 | **0.840** |
| Heart failure | 0.722 | 0.770 | 0.738 | 0.821 | **0.950** |
| Acute myeloid leukemia | 0.839 | 0.858 | 0.853 | 0.915 | **0.957** |
| Lung neoplasm | 0.827 | 0.835 | 0.855 | 0.876 | **0.892** |
| Melanoma | 0.784 | 0.790 | 0.807 | 0.837 | **0.951** |
| Ovarian neoplasm | 0.882 | 0.884 | 0.909 | 0.923 | **0.949** |
| Pancreatic neoplasm | 0.871 | 0.895 | 0.887 | 0.945 | **0.954** |
| Prostatic neoplasm | 0.823 | 0.854 | 0.841 | 0.882 | **0.928** |
| Stomach neoplasm | 0.779 | 0.787 | 0.797 | 0.821 | **0.859** |
| Urinary bladder neoplasm | 0.821 | 0.850 | 0.845 | 0.897 | **0.898** |
| Average AUC | 0.799 | 0.816 | 0.826 | 0.862 | **0.914** |

We compare SPM with MIDP in *AUC* and *REC*, see Figure 4. The *AUC* value of MIDP corresponding to all diseases is 0.829, whereas the *AUC* value of SPM is 0.882, which improves by 5.3%. Moreover, as shown in Figure 5, within the top 30, the average recall of MIDP and SPM for all fifteen diseases are 43.5% and 49.4%, respectively. The *REC* for SPM is higher by 5.9% than that for MIDP. SPM produces superior results by considering the universal structural feature information.

**Fig. 4.**
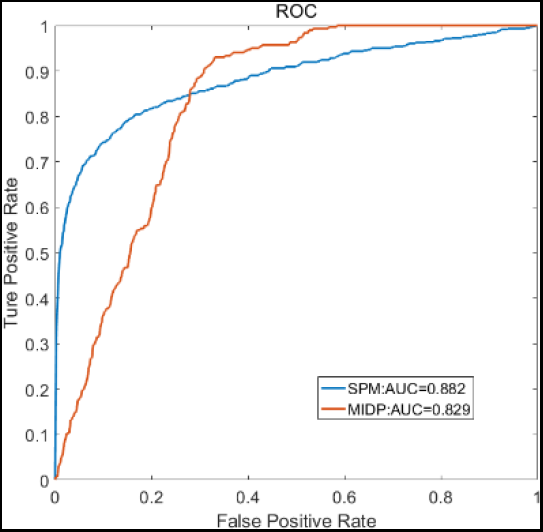
The ROC across all tested diseases

**Fig. 5.**
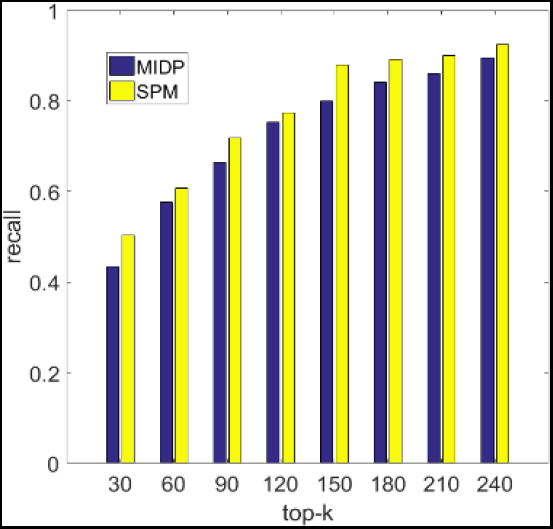
Average recalls across 15 tested diseases

### 3.4 Case study: breast neoplasm

Now we analyze in detail the prediction accuracy on breast neoplasm, and mainly focus on the top-20 miRNA candidates in Table 5. We adopt three ways to evaluate the performance. First, we use the miR2Disease, a database including manually curated miRNAs, which is abnormally regulated in multiple diseases (Jiang, *et al.*, 2009). As shown in Table 5, 6 of 20 candidates are contained in miR2Disease. Secondly, we uses dbDEMC, a database to identify the potential cancer-related miRNAs (Yang, 2010). Table 5 shows that 9 of 20 candidates are included in dbDEMC, indicating that they are upregulated or downregulated in breast neoplasm. Besides, we also find that 8 miRNAs have been supported by literatures, see the candidates labeled with “literature” in Table 5. In general, 16 of 20 candidates can be confirmed by other databases or literatures.

**Table 5.**
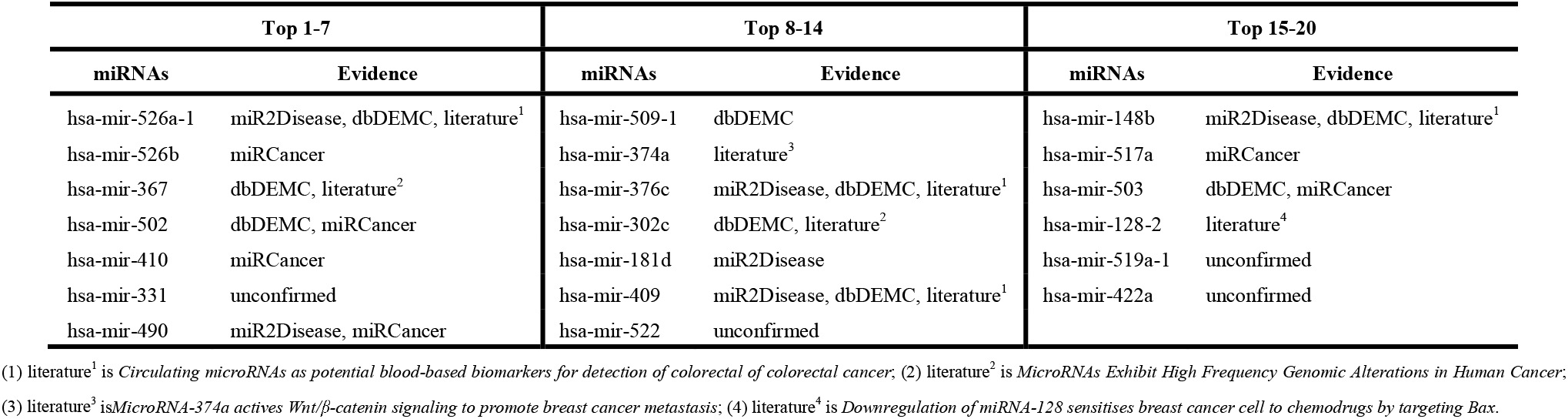
The top-20 breast neoplasm related candidates

| Top 1-7 |  | Top 8-14 |  | Top 15-20 |  |
| --- | --- | --- | --- | --- | --- |
| miRNAs | Evidence | miRNAs | Evidence | miRNAs | Evidence |
| hsa-mir-526a-1 | miR2Disease, dbDEMC, literature<sup>1</sup> | hsa-mir-509-1 | dbDEMC | hsa-mir-148b | miR2Disease, dbDEMC, literature<sup>1</sup> |
| hsa-mir-526b | miRCancer | hsa-mir-374a | literature<sup>3</sup> | hsa-mir-517a | miRCancer |
| hsa-mir-367 | dbDEMC, literature<sup>2</sup> | hsa-mir-376c | miR2Disease, dbDEMC, literature<sup>1</sup> | hsa-mir-503 | dbDEMC, miRCancer |
| hsa-mir-502 | dbDEMC, miRCancer | hsa-mir-302c | dbDEMC, literature<sup>2</sup> | hsa-mir-128-2 | literature<sup>4</sup> |
| hsa-mir-410 | miRCancer | hsa-mir-181d | miR2Disease | hsa-mir-519a-1 | unconfirmed |
| hsa-mir-331 | unconfirmed | hsa-mir-409 | miR2Disease, dbDEMC, literature<sup>1</sup> | hsa-mir-422a | unconfirmed |
| hsa-mir-490 | miR2Disease, miRCancer | hsa-mir-522 | unconfirmed |  |  |

(1) literature<sup>1</sup> is *Circulating microRNAs as potential blood-based biomarkers for detection of colorectal of colorectal cancer*; (2) literature<sup>2</sup> is *MicroRNAs Exhibit High Frequency Genomic Alterations in Human Cancer*; (3) literature<sup>3</sup> is *MicroRNA-374a activates Wnt/ $\beta$ -catenin signaling to promote breast cancer metastasis*; (4) literature<sup>4</sup> is *Downregulation of miRNA-128 sensitises breast cancer cell to chemodrugs by targeting Bax*.

To further manifest the performance of SPM, we use the old version of the human miRNA disease database (HMDDv1.0, before June 16, 2013) as training data to construct the bilayer network and to predict the associations between miRNAs and diseases; then we employee the newly discovered disease miRNAs in HMDDv2.0 to verify the prediction results. In this experiment, we still use breast neoplasm as a case study because the newly discovered breast neoplasm related miRNAs are the most among all the diseases. Figure 6 shows the recalls of new discovered miRNAs of breast neoplasm predicted by SPM and MIDP. In all six groups of investigations, SPM outperforms MIDP. For the top-50 predicted miRNAs, the recall rate of SPM and MIDP are 29.4%, and 17.6%, respectively. The improvement increases if we focus on the top-100 places.

**Fig. 6.**
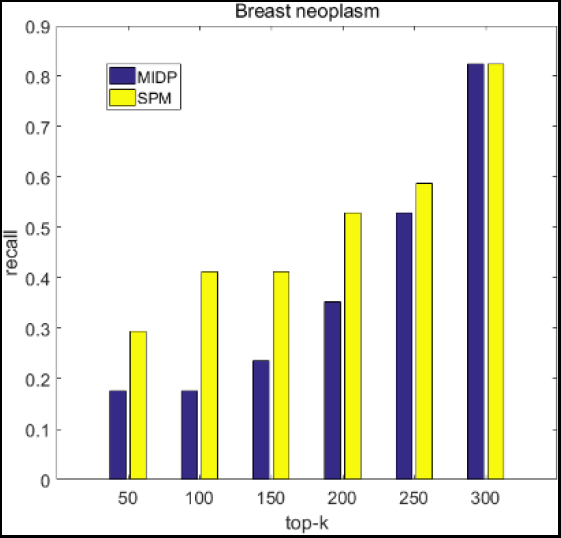
Recalls of newly discovered miRNAs of breast neoplasm

In summary, the results of case studies on breast neoplasm also demonstrate that SPM is a useful tool to identify the potential disease-associated miRNAs.

## 4 Conclusion

We constructed a miRNA–disease bilayer network which consists of three single networks, namely the verified disease-related miRNAs associations, the diseases similarity network and the miRNAs similarity network. We used the structural consistency indicator to quantify the links predictability of this network and found that the miRNA-disease bilayer network is much more predictable than any of the three single networks. Subsequently, we applied the structural perturbation method (SPM) to predict the connections of miRNAs-diseases in bilayer network. For SPM, we finally obtained the perturbed matrix, which is considered as the score matrix between pair of nodes in the network. SPM does not use the similarity of two nodes to make prediction, but recovers the missing links (i.e., unknown information) by perturbing the network with another set of known links.

We compared SPM with RWRMDA, HDMP, RLSMDA, and MIDP. The results demonstrate that SPM is powerful in discovering potential disease-related miRNA candidates. Furthermore, a case study on breast neoplasm was carried out and a good prediction can also be achieved by SPM. We believe that SPM is useful in supplying reliable candidates for further study on the involvement of disease pathogenesis.

